# LAMPP: A benchmark for continuous evaluation of host phenotype prediction from shotgun metagenomic data

**DOI:** 10.1101/2025.06.12.658885

**Authors:** Netta Barak, Haimasree Bhattacharya, Francesco Asnicar, Jaeyun Sung, Nicola Segata, Moran Yassour

**Affiliations:** School of Computer Science and Engineering, The Hebrew University of Jerusalem, Jerusalem 91904, Israel; Center for Interdisciplinary Data Science Research (CIDR) at the Hebrew University of Jerusalem, 91904, Israel; Department of Cellular, Computational and Integrative Biology, University of Trento, 38123 Trento, Italy; Division of Computational Biology, Department of Quantitative Health Sciences, Mayo Clinic, Rochester, MN 55905, USA; Department of Microbiology and Molecular Genetics, IMRIC, Faculty of Medicine, The Hebrew University of Jerusalem, Jerusalem 91121, Israel; Computational Medicine Center, The Hebrew University of Jerusalem, Faculty of Medicine, The Hebrew University of Jerusalem, Jerusalem 91121, Israel

## Abstract

Predicting host phenotypes from shotgun metagenomic data is essential for translating microbiome research into clinical practice)Despite the development of numerous computational tools for this task, researchers often default to traditional machine learning methods such as Random Forest)This hesitancy to adopt newer methods stems from their complexity as well as the lack of standardized evaluations, as most tools are assessed on different datasets and compared against a limited set of methods)Here, we introduce LAMPP, a standardized benchmark for evaluating methods for predicting host phenotypes from gut metagenomic data)LAMPP features a diverse range of prediction tasks and enables consistent, comparative assessments across prediction tools)Our systematic evaluation of existing tools shows that classic machine learning methods (e.g., Random Forest) perform competitively, offering both ease of use and state-of-the-art results)At the same time, it demonstrates that microbiome-based phenotype prediction remains a challenging problem)By providing a consistent platform for ongoing evaluation and access to raw sequencing data, LAMPP motivates the development of novel prediction pipelines from raw sequencing data to phenotype prediction, including novel sample representation and data augmentation strategies)LAMPP is publicly available for ongoing benchmarking at https://lampp.yassourlab.com/.

## Introduction

The human microbiome, consisting of trillions of microorganisms, plays a critical role in various aspects of human health^1,2^)Predicting host phenotypes from microbiome samples is a fundamental task in microbiome research and a foundational step toward translating it into clinical applications, such as disease diagnosis^3–5^, prognosis^6,7^, monitoring^8^, and treatment selection^9^)Recent advances in microbiome research have significantly enhanced the ability to pursue this goal)In particular, the widespread adoption of shotgun metagenomic sequencing has enabled a more comprehensive characterization of microbial communities by capturing genetic material across entire genomes^10^, in contrast to the more targeted 16S rRNA sequencing)These developments, along with improved tools for taxonomic^11,12^ and functional^13^ profiling, have made it increasingly feasible to approach host phenotype prediction as a supervised classification task)In response, a growing number of specialized tools^14–24^ have been developed to tackle this task, many of which leverage deep learning techniques^16–23,25^ — often described as a promising direction for unlocking the full potential of microbiome-based prediction^26,27^.

Despite the proliferation of specialized tools, there is a substantial gap between methodological development and practical implementation)In practice, researchers still largely rely on generating taxonomic profiles and then applying conventional general-purpose machine learning approaches such as Random Forest^28,29^)The limited employment of novel tools stems from two critical challenges)First, tools are often benchmarked using different tasks, datasets, and evaluation metrics, making it challenging for researchers to compare performance or identify methods best suited to their data)In the absence of standardized evaluations, it is both reasonable and practical for researchers to rely on simple and familiar approaches)Second, major usability problems make it difficult to adopt novel tools)This includes heavy computational demands (e.g., reliance on GPUs or long runtime on CPUs), outdated and poorly maintained code, insufficient documentation, and considerable effort and expertise required to adapt existing implementations to a standard train-and-inference scheme.

Independent benchmarks have been foundational to the AI revolution, as demonstrated by the transformative impact of efforts such as ImageNet^30^ for computer vision and CASP^31,32^ for protein folding)These frameworks provide standardized datasets and rigorous evaluation metrics, enabling the unbiased comparisons necessary to catalyze the development and adoption of novel methodologies)While previous independent benchmarks for metagenomic host phenotype prediction provided initial insights^15,33–35^, their coverage remained limited by their static nature)By prioritizing cross-validation-based statistical rigor, these studies often sacrificed scalability and flexibility; consequently, they represent only a snapshot of available tools and cannot integrate new methodologies post-publication)-)Live benchmarks address these limitations and offer a dynamic alternative)By facilitating continuous, unbiased evaluation, they move beyond a pre-selected set of tools, shifting the responsibility to developers to demonstrate their performance within a standardized framework)While this model is well-established in other computer science disciplines^36^, live benchmarking was only recently introduced to the microbiome community via the CAMI portal^37^ for benchmarking assembly, binning, and taxonomic profiling)Beyond performance tracking, such benchmarks establish minimum standards for method developers, incentivizing the maintenance of accessible code repositories, particularly for high-performing approaches)By directly addressing the current challenges in microbiome-based host phenotype prediction, comprehensive live benchmarks not only highlight existing limitations but also drive the development of more robust, user-friendly methods tailored to real-world challenges.

Here, we present LAMPP (Live Assessment of Metagenomics-based tools for host Phenotype Prediction), a standardized, comprehensive, and continuously updated benchmark for host phenotype prediction methods)LAMPP provides a unified framework for evaluating prediction methods over a diverse set of host phenotype binary classification tasks based on gut metagenomic data)These tasks vary in phenotype targets, training set composition, and training-test similarity, reflecting real-world prediction scenarios)LAMPP is open to the scientific community for evaluating their tools and submitting their prediction results, keeping the benchmark current, dynamic, and relevant to the evolving state of the field.

## Results

### Prediction tasks design

In developing LAMPP, we aimed to provide a platform for consistent comparison of host phenotype prediction tools across diverse scenarios, helping users identify methods that are either robust across various scenarios or well-suited to tasks similar to their own (**Figure 1**)a))We also aimed to highlight current challenges in the field and showcase the limitations of existing tools, thereby pointing out opportunities for future methodological improvements.

**Fig)1:**
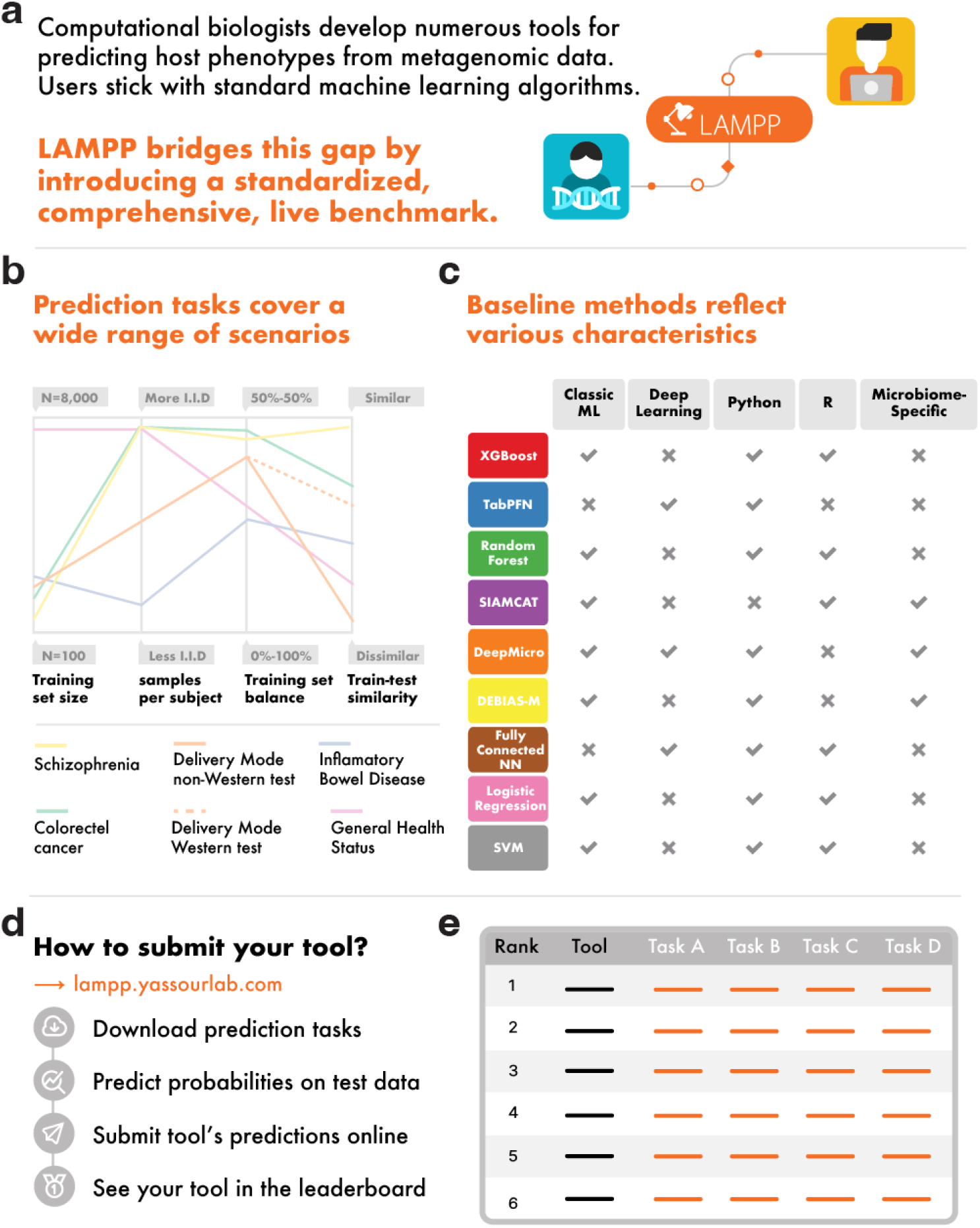
Overview of LAMPP’s benchmarking platform for host phenotype prediction. **a**, LAMPP was developed to enable the ongoing evaluation of tools for predicting host phenotypes from shotgun metagenomics)**b**, To facilitate the comparison of different tools across diverse scenarios, the tasks are designed to vary in training set size, number of samples per subject, class balance, and the similarity between training and test sets)**c**, The selected baseline prediction tools capture diverse methodological and practical aspects)**d**, Developers can submit their tools to be evaluated using LAMPP’s website)**e**, The performance of all evaluated tools across all tasks can be compared through LAMPP’s online leaderboard.

With these aims in mind, we designed LAMPP’s prediction tasks to span a range of real-world gut-metagenomic-based classification scenarios)Each task is framed as a binary classification problem, with a labeled training set and an unlabeled test set, both derived from publicly available data (**Methods**))The tasks vary in the predicted phenotype and the design of their training and test sets, reflecting the diversity and complexity of metagenomic data (**Figure 1**)b).

The training sets were designed to reflect common challenges in the field)They vary in dataset size, class balance, sample dependence (mostly introduced by longitudinal sampling), and batch effects (mostly caused by combining datasets that differ in their protocols for data collection and processing).

Test sets were designed to evaluate generalization under varying degrees of resemblance to the training set)One task (schizophrenia) uses test data from the same cohort as the training set, representing maximal resemblance between the sets)In the remaining tasks, test sets are drawn from distinct cohorts — ranging from similar populations to those that differ substantially in geography, lifestyle, and disease context — enabling evaluation across a spectrum of generalization scenarios)To ensure reliable performance metrics, test sets include only one sample per individual and maintain a substantial representation (above 25%) of both classes.

Here is the description of all current prediction tasks included in LAMPP, which could be extended in the future to account for future challenges:

#### Colorectal Cancer (CRC) Task

This task aims to reflect a common use case in which researchers demonstrate the application of host phenotype prediction for disease diagnosis)The training set is balanced, contains one sample per individual, and comprises 983 samples from 498 people with colorectal cancer (CRC) and 485 healthy controls, pooled from eight studies in the USA^38^, Japan^39^, France^5^, Italy^40^, Austria^41^, China^42^, and India^43^)The test set includes 125 samples from people with CRC and healthy controls collected independently in Germany^3^.

#### Inflammatory Bowel Disease (IBD) Task

Like the Colorectal Cancer task, this task was also designed as a microbiome-based disease diagnosis scenario, but with key differences that make it more challenging)In addition to predicting a different phenotype, the IBD training set is slightly imbalanced (roughly 3:1 IBD vs)control ratio), increasing the risk of models favoring the majority class)In addition, the training set involves longitudinal sampling, namely, it includes multiple samples that were collected from the same individual over time)Longitudinal sampling is a common practice in microbiome research as it allows tracking the dynamics of the microbiome composition)However, in the scenario of training ML algorithms, it introduces dependencies between samples and violates the assumption of independent and identically distributed samples (*i*.*i*.*d*.) underlying many predictive models)The training set comprises 1,886 longitudinal samples from 123 people with IBD and 40 healthy controls across two USA cohorts^44,45^ (ages 6–76, median 21))The test set includes 506 samples (one sample per person, ages 7–70, median 47) drawn from three studies in Spain^46,47^, Denmark^46,47^, the United Kingdom^48^, and China^46^.

#### Delivery Mode Tasks (with either Western [DMW] or non-Western [DMNW] test set)

Developing tools that can generalize across populations, typically from well-studied to understudied populations, is a key challenge in human microbiome research)To highlight this challenge, we developed a pair of delivery mode prediction tasks that share a training set but differ in their test set)The training set comprised 1,461 longitudinal samples (see comment on longitudinal sampling in the **IBD Task** description) from 461 vaginally delivered and 290 cesarean-delivered healthy term infants (ages 0 to 180 days) in the United Kingdom^49^, Sweden^50^, Finland^51^, Estonia^51^, and Russia^51^)The western test set includes 101 samples from infants aged 0 to 90 days in Finland^52^, Luxembourg^53^, and the USA^54^, while the non-western test set includes 221 samples from infants aged 0 to 180 days in Bangladesh^55^)When designing this pair of tasks, we expect the DMNW to be more challenging, due to the differences between the populations captured in the training and test sets.

#### General Health Status (GHS) Task

In recent years, two complementary approaches have emerged for understanding microbiome patterns associated with health and disease: one seeks to define the characteristics of a healthy microbiome^56–58^ and identify deviations as potential indicators of illness, while the other focuses on identifying common microbial signatures shared across various disease states^59,60^)From a machine learning standpoint, both approaches present similar opportunities and challenges)A major opportunity lies in addressing the typically small sample sizes of microbiome studies — especially given the high dimensionality and complexity of microbiome data)By combining datasets, researchers can compose large training sets, which are generally more effective for developing robust predictive models)However, this approach also brings challenges: (1) training on large-scale data demands efficient computational methods or significant computing resources; (2) the combination of multiple cohorts collected and processed using different experimental protocols introduces noise and batch effects; and finally, (3) *general health* is a complex phenotype, involving multiple or overlapping disease conditions, which adds further difficulty to model development and interpretation.

To highlight this challenge, we developed a task based on the training and test sets used in the Gut Microbiome Wellness Index 2 (GMWI2)^8^)The slightly imbalanced training set consists of 8,069 samples collected from 5,547 healthy individuals and 2,522 non-healthy individuals combined from 54 studies across 26 countries across six continents)Non-healthy individuals were diagnosed with one of 11 diseases: colorectal cancer, Crohn’s disease, ulcerative colitis, type 2 diabetes, atherosclerotic cardiovascular disease, liver cirrhosis, metabolic dysfunction-associated steatotic liver disease, Graves’ disease, multiple sclerosis, rheumatoid arthritis, and ankylosing spondylitis)The test set consists of 1,140 samples from healthy individuals and people diagnosed with pancreatic cancer, ankylosing spondylitis, or Parkinson’s disease from six cohorts collected in Tanzania, Luxembourg, Switzerland, China, Japan, and the USA^8^.

#### Schizophrenia Task (SCZ)

This task represents a scenario where researchers aim to predict a host phenotype for which no additional metagenomic datasets are available, or where they deliberately choose to train and test their model on the same cohort for other reasons (for example, when their cohort is unique in some aspect that is difficult to account for))In the Schizophrenia task, we use a dataset of 171 samples from treatment-naïve people with schizophrenia and healthy controls in China^61^, randomly split into training and test sets)The training set is balanced and includes 119 samples, comprising 63 people with schizophrenia and 56 healthy controls, and the test set contains the remaining 52 samples.

### Baseline prediction tools

LAMPP was designed to provide a common platform for the ongoing evaluation of host phenotype prediction tools as they emerge)To ensure informative comparisons, we chose to include a set of baseline methods representing the current state of the art)We selected baseline tools that implement various strategies, including differences in feature selection, data normalization, sample representation, programming languages, and modeling approaches, ranging from traditional machine learning to deep learning.

The baseline tools include Random Forest^62^, XGBoost^63^, Logistic Regression^64^, SVM^65^, SIAMCAT^15^, DeepMicro^16^, DEBIAS-M^66^, TabPFN^67,68^ and a Fully Connected Neural Network^69^ (**Figure 1**)c; **Methods**))We also attempted to include PopPhy-CNN^70^; however, despite our continuous attempts, we were not able to run it to completion, thus it was not included in our evaluation)Most baselines are popular off-the-shelf implementations of machine learning algorithms, while SIAMCAT, DEBIAS-M, and DeepMicro are tools tailored specifically for metagenomics data)These tools incorporate additional preprocessing steps (including feature selection, normalization, and dimensionality reduction) prior to applying a machine learning classifier (**Methods**))We ran a parameter search for each tool using cross-validation on the training set (**Methods; Supplementary Information**), and presented here the optimized-parameter version (**Figure 2**), alongside the default option (**Supplementary Information**).

**Fig)2:**
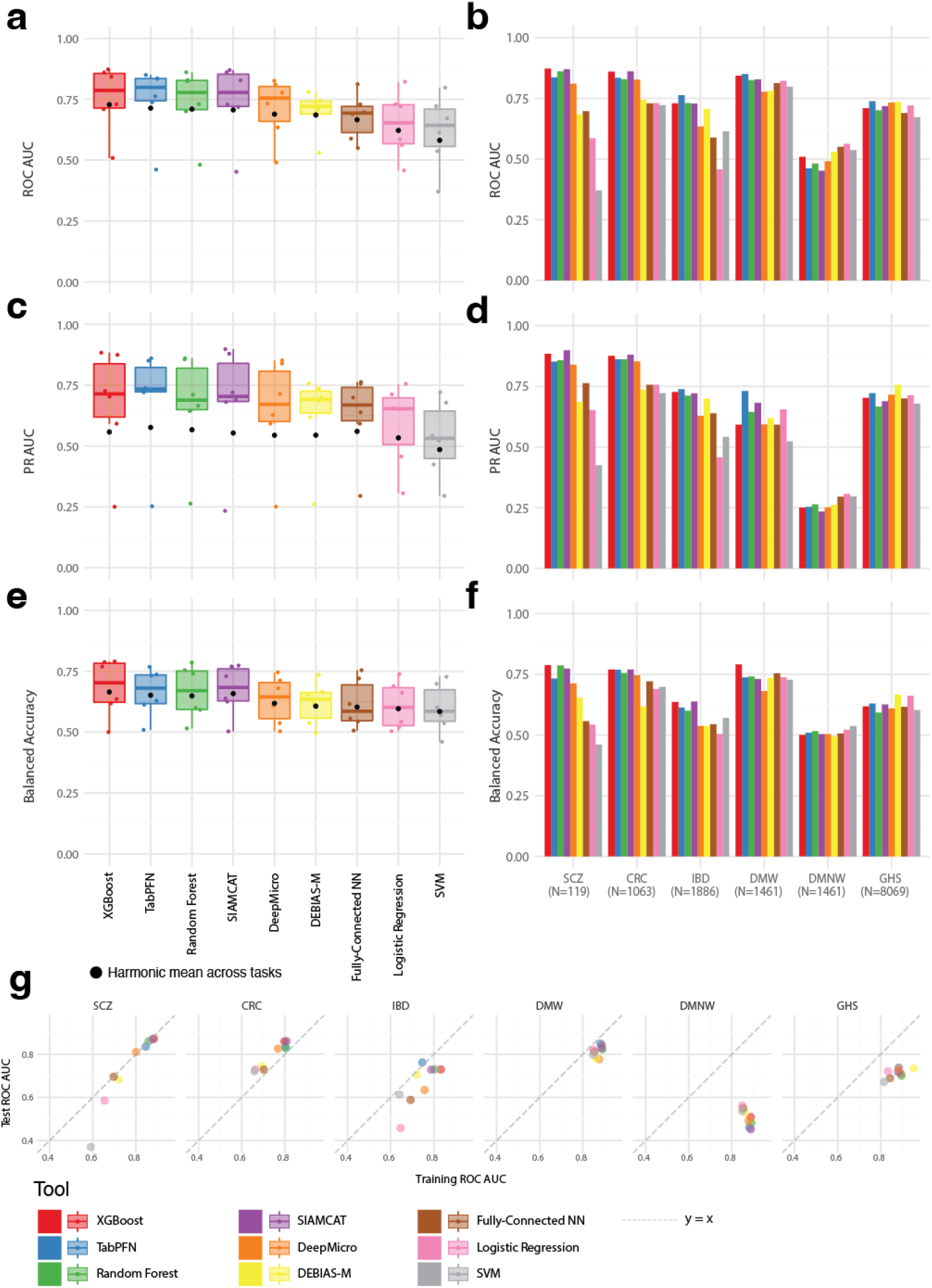
Performance of baseline tools with optimized parameters. **a-f**, A comparison of the performance of baseline prediction tools over the Schizophrenia (SCZ), Colorectal Cancer (CRC), Delivery Mode with Western (DMW) and Non-Western (DMNW) test sets, Inflammatory Bowel disease (IBD), and General Health Status (GHS) tasks)ROC AUC (**a, b**), Precision-Recall (PR) AUC (**c, d**), and balanced accuracy (**e, f**) of baseline tools (color), per task (**b, d, f**), and overall (**a**,**c**,**e**))Black points mark the harmonic mean across tasks)**g**, Comparing training (x-axis) and test (y-axis) set ROC AUC, using the optimized parameters on the training datasets, across tools (colors) and tasks (panels); Dashed line marks y=x)The training set ROC AUC is averaged over training cross-validation folds)Overfitting is observed here when the training ROC AUC is much higher than the test ROC AUC.

### Performance of baseline tools

To evaluate the performance of all tools across all tasks, we first generated the species-level taxonomic profiles for all samples (**Methods**))For every task, we trained all tools on the training set using both the default and the optimized parameters and then predicted class probabilities on the test set)We evaluated the tools’ performance using multiple metrics: area under the receiver operating characteristic curve (ROC AUC), area under the precision-recall curve (PR AUC), balanced accuracy, precision, recall, and F1 score (**Figure 2, Figure S1-S4; Methods**))Finally, to favor methods that demonstrate robustness and consistency across prediction scenarios over those that excel in some but perform poorly in others, we chose the harmonic mean across all tasks (rather than the arithmetic mean, **Methods**) as our overall score (**Tables S2-S8**).

We considered the overall performance of the runs with optimized parameters based on ROC AUC, PR AUC, and balanced accuracy (**Figure 2**)a-f), and found the top-performing tools to be XGBoost (top ROC AUC = 0.728, balanced accuracy = 0.665) and TabPFN (top PR AUC = 0.577))The lowest ROC AUC, PR AUC, and balanced accuracy were achieved by SVM (ROC AUC = 0.581; PR AUC = 0.486; balanced accuracy = 0.585; **Figure 2**)a, c, and e, **Tables S2-S4**).

Next, to further explore the various challenges that each task presents, as well as evaluate the potential overfitting of some tools, we examined the differences in performance on the training and test data sets)We compared the ROC AUC on the test dataset to the ROC AUC of the optimized runs, averaged over the cross-validation folds (hereafter *training ROC AUC*; **Figure 2.** g))This comparison suggested a few patterns)For the SCZ, CRC, and DMW tasks, test ROC AUCs were similar to the corresponding training values (ranged from an increase of of 6.55% to a decrease of 6.369% between the training ROC AUC and test ROC AUC), suggesting overall no cases of overfitting)In contrast, for the IBD and GHS task, training ROC AUC and test ROC AUC decreased in 10.424% and 18.497%, respectively, indicating some overfitting)Additionally, for the DMNW, all tools achieved training ROC AUCs between 0.848 and 0.888 while test ROC AUCs hovered around 0.5 (**Figure 2**)b, i.e., no better than random guessing), suggesting no clear generalization of the microbiome signature for this task)One possible explanation for this lack of generalization is a mismatch between the cohorts used for training and testing (see **Delivery Mode Tasks**))Together, these results illustrate that the magnitude of train–test domain shift varies substantially across tasks, making some considerably more challenging than others.

### Runtime and usability of baseline tools

While prediction accuracy is a critical factor when selecting a tool for host phenotype prediction, it is not the only consideration)In practice, widespread adoption of a tool also depends on factors such as runtime, computational efficiency, active maintenance, clear documentation, and a user-friendly interface for training and inference (as tool developers usually demonstrate the performance of their tools using cross-validation).

To assess runtime performance, we ran all tools using identical computational resources (**Methods**) and measured the total runtime of each parameter search configuration (**Table 1; Figure S5**))As expected, the runtime increased with the size of the training dataset)SIAMCAT and DEBIAS-M took the longest to train, reaching an average training time of 100.8 minutes and 74.2 minutes, respectively, for each parameter configuration tested)The slow run time of these tools was mostly evident in the General Health Status task (N=8,069), where the average runtime per tested configuration was 535 minutes for SIAMCAT and 317 minutes for DEBIAS-M)This is while training a single configuration for Random Forest, XGBoost, or the Fully Connected NN took an average of less than two minutes for all the tasks.

**Table 1:**
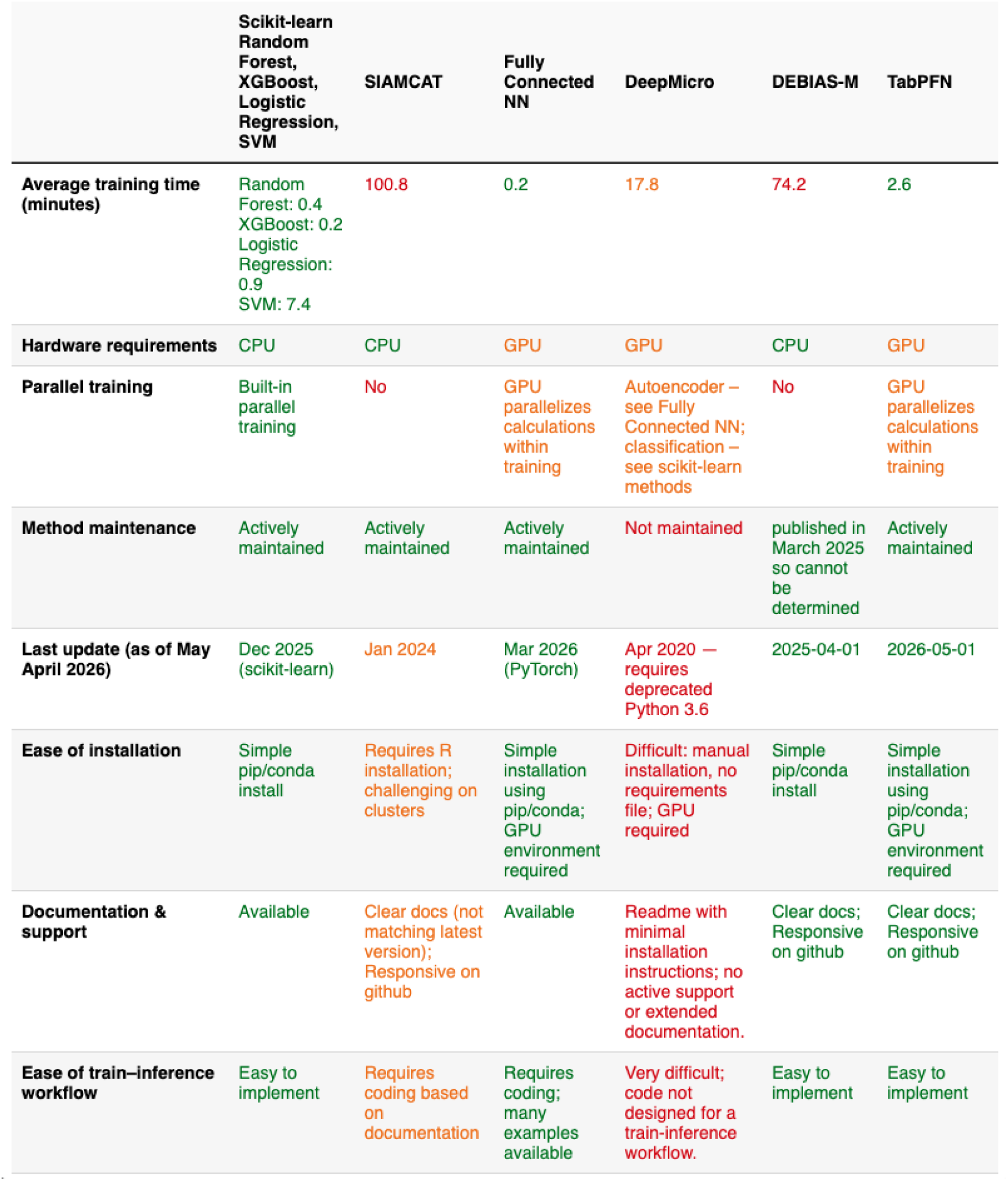
Usability evaluation of baseline prediction tools. The usability of prediction tools was evaluated across multiple criteria)Random Forest, XGBoost, Logistic Regression, and SVM are presented together as they were implemented using the same software package and share a similar user experience)Colors indicate the ease of use for each criterion assessed: green = easy, orange = intermediate, red = difficult.

|  | Scikit-learn<br>Random<br>Forest,<br>XGBoost,<br>Logistic<br>Regression,<br>SVM | SIAMCAT | Fully<br>Connected<br>NN | DeepMicro | DEBIAS-M | TabPFN |
| --- | --- | --- | --- | --- | --- | --- |
| Average training time (minutes) | Random Forest: 0.4<br>XGBoost: 0.2<br>Logistic Regression: 0.9<br>SVM: 7.4 | 100.8 | 0.2 | 17.8 | 74.2 | 2.6 |
| Hardware requirements | CPU | CPU | GPU | GPU | CPU | GPU |
| Parallel training | Built-in parallel training | No | GPU parallelizes calculations within training | Autoencoder – see Fully Connected NN; classification – see scikit-learn methods | No | GPU parallelizes calculations within training |
| Method maintenance | Actively maintained | Actively maintained | Actively maintained | Not maintained | published in March 2025 so cannot be determined | Actively maintained |
| Last update (as of May April 2026) | Dec 2025 (scikit-learn) | Jan 2024 | Mar 2026 (PyTorch) | Apr 2020 — requires deprecated Python 3.6 | 2025-04-01 | 2026-05-01 |
| Ease of installation | Simple pip/conda install | Requires R installation; challenging on clusters | Simple installation using pip/conda; GPU environment required | Difficult: manual installation, no requirements file; GPU required | Simple pip/conda install | Simple installation using pip/conda; GPU environment required |
| Documentation & support | Available | Clear docs (not matching latest version); Responsive on github | Available | Readme with minimal installation instructions; no active support or extended documentation. | Clear docs; Responsive on github | Clear docs; Responsive on github |
| Ease of train–inference workflow | Easy to implement | Requires coding based on documentation | Requires coding; many examples available | Very difficult; code not designed for a train-inference workflow. | Easy to implement | Easy to implement |

To evaluate usability, we defined several criteria: hardware requirements, parallel training capabilities, maintenance status, ease of installation, availability of tutorials and user support, and the simplicity of executing a standard training and inference scheme)Each method was assessed against all these criteria (**Table 1**).

The majority of tools we tested were available as standard Python and R packages, supported by proper documentation, code examples, and Q&A forums)In contrast, microbiome-specific deep learning tools like DeepMicro and PopPhy-CNN, primarily available via GitHub repositories, were the most difficult to use)These tools suffered from insufficient maintenance, outdated or missing dependencies, incomplete installation instructions, and often required major code modifications to support simple training and inference workflows.

### LAMPP’s continuous evaluation platform

LAMPP’s primary objective is to serve as a platform for the ongoing evaluation of emerging prediction tools (**Figure 1**)d))To make sure that this effort does not remain a static table of the currently selected tools, and also enables the evaluation of newly developed tools, we made LAMPP publicly available at https://lampp.yassourlab.com)The prediction tasks’ data are available both as raw metagenomic sequencing data and as species-level taxonomic profiles (MetaPhlAn^12^, **Methods**), together with subject IDs, enabling developers to adjust for longitudinal sampling if they choose to do so)Developers can download the tasks data, apply their tools, and submit the predicted class probabilities)These submissions are automatically evaluated, with results displayed on a public leaderboard that enables transparent comparison across tools (**Figure 1**)e).

As new metagenomic datasets become available, they offer opportunities to explore novel challenges and improve predictive modeling)To support this, LAMPP is built to accommodate new tasks over time)We encourage researchers to contribute by suggesting new tasks or sharing their soon-to-be-published datasets as temporarily hidden test sets)Tool developers will be notified and invited to make new submissions when new tasks are added.

## Discussion

Here, we introduce LAMPP, a standardized benchmark for the continuous evaluation of tools for predicting host phenotypes from gut metagenomic data)LAMPP’s binary classification tasks cover a diverse range of prediction scenarios and host phenotypes, reflecting the common practices and key challenges in microbiome research)Our evaluation of baseline prediction tools reinforces existing trends: although novel methods such as TabPFN perform competitively, tree-based approaches, such as XGBoost and Random Forest, consistently achieve strong performance while maintaining computational efficiency and user accessibility.

Despite these strengths of Random Forest, our analysis highlights that host phenotype prediction from metagenomic data remains a difficult task — particularly when generalizing to understudied populations)Moreover, we observed only modest performance differences across the evaluated tools)This limited performance variation may stem from multiple factors: (1) the benchmarked tools lack sufficient methodological diversity; (2) taxonomic profiles may not capture the most informative features for these tasks; or (3) the data may lack biological signal, limiting the predictive potential regardless of the method used)To further understand the contribution of each factor, we encourage the development of innovative workflows that view host phenotype prediction from end to end and include aspects such as feature extraction, data augmentation, feature selection, and normalization)To facilitate additional sample representations that go beyond the taxonomic profiles used in our analysis, LAMPP also provides the raw sequencing data for all samples used in the benchmark.

One limitation of LAMPP’s ongoing live evaluation framework is the lack of runtime and usability assessments, as each developer runs their tools on their own infrastructure (to make the submission of new tools to LAMPP as easy as possible))Therefore, we cannot directly evaluate computational efficiency or ease of use, and we choose not to rely on self-reported performance metrics of these aspects)Despite this, our analysis shows that classic machine learning methods not only offer strong predictive performance but also come with efficient, user-friendly implementations)This establishes a high standard for the developers of new tools, emphasizing the importance of maintainability, usability, and efficiency)Especially for the top-ranking tools, we expect the users to demand better usability all around.

In designing this benchmark, we faced a tradeoff: whether to offer diverse, standardized tasks based on real-world public data (rather than simulated data) or restrict the benchmark to limited, private test sets to reduce the risk of overfitting)We chose the former, placing trust in the scientific integrity of tool developers)By submitting to LAMPP, participants affirm that their predictions represent genuine generalization to unseen data)That said, we invite researchers with soon-to-be-published data to collaborate with us in developing new benchmark tasks with a temporarily private test set)These occasional challenges will be advertised to all previous submitters of LAMPP, to enable a comprehensive evaluation of their tools on new and unseen test data.

To conclude, LAMPP represents an important first step in bridging the gap between tool developers and researchers seeking to apply host phenotype prediction in microbiome studies)By providing a transparent, evolving, and community-driven benchmark, LAMPP helps identify the most suitable tools for diverse prediction tasks, encourages methodological innovation, and promotes the development of user-friendly, reproducible software)By making raw sequencing data accessible, LAMPP sets the foundations for comparing future deep learning approaches involving novel sample representations and data augmentation)We invite the entire microbiome research community to make use of this benchmark and continuously contribute additional tools and datasets to keep this resource useful, relevant, and up to date.

## Methods

### Tasks Data

Species-level Taxonomic profiles were generated using MetaPhlAn4^12^ with the *mpa_vJan21_CHOCOPhlAnSGB_202103* database for all tasks, except for the General Health Status (GHS) task)For this task, MetaPhlAn3^13^ with the *mpa_v30_CHOCOPhlAn_201901* database was used to maintain consistency with the original Gut Microbiome Wellness Index 2^8^ (GMWI2) study.

Samples from individuals reported as diagnosed with diseases other than the one being tested were excluded)For the Inflammatory Bowel Disease task, samples from individuals under 5 years of age were excluded)When age was reported categorically, individuals outside the school-age, adult, or senior categories were excluded)For the delivery mode tasks, only full-term, healthy infants aged between 0 and 180 days were included.

### Baseline prediction tools

Unless stated otherwise, the tools were used through their Python implementations available in Scikit-Learn v1.6.0 ^71^.

*Random Forest*^*62*^ is an ensemble learning method that builds multiple decision trees during training and outputs the mode of their predictions at inference)Averaging across trees trained on different random subsets of the data and features makes it both robust and interpretable.

*XGBoost*^*63*^ (Extreme Gradient Boosting) is an ensemble method based on gradient-boosted decision trees)It builds trees sequentially, with each new tree aiming to correct the errors of its predecessors by optimizing a loss function using gradient descent.

*Logistic Regression*^*64*^ is a linear classification method that models the probability of a binary outcome using the logistic (sigmoid) function)It estimates coefficients for input features that best separate the classes and is widely used due to its simplicity and interpretability.

*Support Vector Machine* (SVM)^65^ is a supervised learning algorithm that classifies data by finding the optimal hyperplane separating the classes)It does so by maximizing the margin between the closest points of each class.

SIAMCAT v2.10.0 (Statistical Inference of Associations between Microbial Communities And host phenoTypes)^15^ is a machine learning toolbox specifically designed for microbiome data analysis)It integrates feature selection, cross-validation, phenotype prediction (using logistic regression with various regularization techniques or Random Forest), and performance evaluation tailored to microbiome-specific challenges.*DEBIAS-M (domain adaptation with phenotype estimation and batch integration across studies of the microbiome)*^*66*^ is a computational framework for inference and correction of processing bias in metagenomic data)It learns taxon-specific bias-correction factors for each batch to simultaneously minimize batch effects and maximize cross-study associations with the predicted phenotype)Specifically,

*OnlineDebiasMClassifier* was used in the analysis presented in this work, as it does not require test data to learn the bias-correction factors.

TabPFN (Tabular Prior-data Fitted Network)^67^ is a foundation deep-learning model designed specifically for tabular data)Having been pre-trained on millions of synthetic datasets, it utilizes **in-context learning** to perform classification and regression)This approach allows the model to process a training set as input during inference, enabling instant predictions without modifying the model’s underlying weights.

*A fully connected Neural Network* (Fully Connected NN or FCNN)^69^ is a basic feedforward neural network architecture, consisting of layers of neurons where each neuron is connected to all neurons in adjacent layers)*DeepMicro*^*16*^ is a deep learning framework designed for microbiome-based classification, aiming to capture complex, nonlinear patterns in the data)It uses autoencoders to learn compact representations of high-dimensional microbiome data, which are then passed onto a classifier (*Random Forest, SVM, or FCNN*))To be evaluated on LAMPP’s tasks, DeepMicro was adapted for a train-and-inference workflow (**Code Availability**).

### Parameters optimization

Each tool was run using default parameters or the top-performing parameter configurations as described in the original publication, if no default parameters existed (Methods S1))In addition, each tool was tuned using a parameter grid of up to roughly 100 candidate configurations (Methods S1))The best configuration was selected based on the average ROC AUC of 5-fold cross-validation (Methods S2).

### Evaluation metrics

For the description of the metrics used in this study, we will use the following notation: `TP` `=` `True` `Positives,` `FP` `=` `False` `Positives,` `TN` `=` `True` `Negatives,` and `FN` `=` `False` `Negatives`.

*Precision* (`TP/(TP+FP)`) measures how many of the predicted positive cases are positive)*Recall* (`TP/(TP+FN)`; also known as *sensitivity, true positive rate*) measures how many positive cases were correctly identified)*F1 score (*`2` `x` `(Precision` `x` `Recall)/(Precision` *+* `Recall)`) is the harmonic mean of precision and recall, balancing both concerns.

*The area under the receiver operating characteristic curve* (ROC AUC), measures the classifier’s ability to discriminate between classes across all classification thresholds)It is computed by plotting the true positive rate (`TPR=TP/` `(TP+FN)`; also known as recall) against the false positive rate (`FPR=FP/(FP+TN)`) across different thresholds and calculating the area under the curve)An ROC AUC closer to 1.0 indicates stronger separability and better overall classification performance.

The area under the precision-recall curve (PR AUC) summarizes the trade-off between precision (`TP/(TP+FP)`) and recall (`TP/(TP+FN)`) across thresholds, and is particularly useful for imbalanced datasets)Like ROC AUC, it is computed as the area under the curve and higher values (closer to 1) indicate better performance.

Balanced accuracy is an adaptation of the standard accuracy metric)Balanced accuracy for class imbalance by averaging the recall of each class)For binary classification, it is defined as `(TPR+TNR)/2`, where `TPR=Recall=TP/(TP+FN)` and `TNR=Specificity=TN/(TN+FP)`.

For each tool and each metric, the overall score was defined using the harmonic mean of the metric across all tasks)The harmonic mean is defined as *HM*(*x_1_*, *x*_*2*_,..., *x*_*n*_ *) = n/(1/x*_*1*_ *+ 1/x*_*2*_ + ..)+ *1/x*_*n*_), and therefore, if the metric’s value for one or more tasks is 0, the overall score will be undefined)To avoid this, 0 values were replaced with 0.001 for the calculation of the overall score.

### Runtime evaluation

Runtime was evaluated by executing all tools on a machine equipped with an NVIDIA RTX6000 (24GB) GPU and eight CPU cores)Among the evaluated methods, only DeepMicro and FCNN were designed to leverage GPU acceleration)DEBIAS-M is based on the PyTorch^72^ software package that can utilize GPUs to significantly reduce runtime; however, it currently does not officially support the use of GPUs)All CPU-based tools, except SIAMCAT and DEBIAS-M, were designed to utilize multiple CPU cores for parallel processing.

## Data availability

The training and test sample accessions for all tasks, along with their corresponding MetaPhlAn species-level taxonomic profiles, are publicly available at: https://lampp.yassourlab.com/tasks/)A complete list of datasets included in the benchmark is provided in **Table S3**.

The dataset used for the General Health Status (GHS) task is based on publicly available data and was curated by the authors of the GMWI2 study^8^; see the references and Data Availability section of their publication for details.

Sequencing data for the delivery mode task with a non-Western test set were obtained from Vatanen et al.^55^ and are available in the NCBI Sequence Read Archive under BioProject accession PRJNA806984)Metadata on delivery mode for these samples is available upon request, as noted in the Data and Code Availability section of their publication)All remaining datasets used in LAMPP’s tasks are accessible via the CuratedMetagenomicData^73^ resource.

## Code availability

The code used for training and inference with all tools is publicly available at https://github.com/yassourlab/LAMPP-benchmark.

## Acknowledgments

We thank the developers and maintainers of CuratedMetagenomicData for enabling us and others access to large, curated metagenomic datasets in a standardized manner)We also thank the authors of the original studies used in LAMPP’s tasks for contributing their data to the scientific community)We further thank Levi Waldron, A)Murat Eren, Gabriel Stanovsky, and Michael Hassid for helpful discussions and suggestions throughout the course of this work.

## Author Contributions

M.Y)and N.B)conceptualized the study, designed the benchmark tasks, and selected the baseline tools)F.A., J.S., and N.S)provided the preprocessed data for the benchmark)H.B)and N.B)designed and implemented the website)N.B)performed the analyses)M.Y)and N.B)wrote the manuscript, with contributions from F.A., J.S., and N.S.

## References

1. Gilbert, J)A. et al. Current understanding of the human microbiome)Nat)Med)24, 392–400 (2018).

2. Clemente, J)C., Ursell, L)K., Parfrey, L)W. & Knight, R. The impact of the gut microbiota on human health: an integrative view)Cell 148, 1258–1270 (2012).

3. Wirbel, J. et al. Meta-analysis of fecal metagenomes reveals global microbial signatures that are specific for colorectal cancer)Nat)Med)25, 679–689 (2019).

4. Knights, D., Parfrey, L)W., Zaneveld, J., Lozupone, C. & Knight, R. Human-associated microbial signatures: examining their predictive value)Cell Host Microbe 10, 292–296 (2011).

5. Zeller, G. et al. Potential of fecal microbiota for early-stage detection of colorectal cancer)Mol)Syst)Biol)10, 766 (2014).

6. Ferreiro, A)L. et al. Gut microbiome composition may be an indicator of preclinical Alzheimer’s disease)Sci)Transl)Med)15, eabo2984 (2023).

7. Leustean, A)M. et al. Implications of the Intestinal Microbiota in Diagnosing the Progression of Diabetes and the Presence of Cardiovascular Complications)Journal of Diabetes Research 2018, 1–9 (2018).

8. Chang, D. et al. Gut Microbiome Wellness Index 2 enhances health status prediction from gut microbiome taxonomic profiles)Nat)Commun)15, 7447 (2024).

9. d’Humières, C. et al. The potential role of clinical metagenomics in infectious diseases: Therapeutic perspectives)Drugs 81, 1453–1466 (2021).

10. Quince, C., Walker, A)W., Simpson, J)T., Loman, N)J. & Segata, N. Shotgun metagenomics, from sampling to analysis)Nat)Biotechnol)35, 833–844 (2017).

11. Wood, D)E., Lu, J. & Langmead, B. Improved metagenomic analysis with Kraken 2)Genome Biol)20, 257 (2019).

12. Blanco-Míguez, A. et al. Extending and improving metagenomic taxonomic profiling with uncharacterized species using MetaPhlAn 4)Nat)Biotechnol)41, 1633–1644 (2023).

13. Beghini, F. et al. Integrating taxonomic, functional, and strain-level profiling of diverse microbial communities with bioBakery 3)Elife 10, e65088 (2021).

14. Nguyen, T)H., Prifti, E., Chevaleyre, Y., Sokolovska, N. & Zucker, J.-D. Disease Classification in Metagenomics with 2D Embeddings and Deep Learning)arXiv [cs.CV] (2018).

15. Wirbel, J. et al. Microbiome meta-analysis and cross-disease comparison enabled by the SIAMCAT machine learning toolbox)Genome Biol)22, 93 (2021).

16. Oh, M. & Zhang, L. DeepMicro: deep representation learning for disease prediction based on microbiome data)Sci)Rep)10, 6026 (2020).

17. Liu, B., Wei, Y., Zhang, Y. & Yang, Q. Deep Neural Networks for High Dimension, Low Sample Size Data)2287–2293 (2017).

18. Danilevsky, A. & Shomron, N. Deep Learning Applied on Next Generation Sequencing Data Analysis)in Deep Sequencing Data Analysis (ed)Shomron, N.) 169–182 (Springer US, New York, NY, 2021).

19. Gadupudi, A. et al. A deep learning framework for human disease prediction using microbiome data)in 2024 International Conference on Integrated Circuits and Communication Systems (ICICACS) 1–6 (IEEE, 2024).

20. Fioravanti, D. et al. Phylogenetic convolutional neural networks in metagenomics)BMC Bioinformatics 19, 49 (2018).

21. Lo, C. & Marculescu, R. MetaNN: accurate classification of host phenotypes from metagenomic data using neural networks)BMC Bioinformatics 20, 314 (2019).

22. Tanaseichuk, O., Borneman, J. & Jiang, T. Phylogeny-based classification of microbial communities)Bioinformatics 30, 449–456 (2014).

23. Sharma, D., Paterson, A)D. & Xu, W. TaxoNN: ensemble of neural networks on stratified microbiome data for disease prediction)Bioinformatics 36, 4544–4550 (2020).

24. Prifti, E. et al. Interpretable and accurate prediction models for metagenomics data)Gigascience 9, giaa010 (2020).

25. Roy, G., Prifti, E., Belda, E. & Zucker, J.-D. Deep learning methods in metagenomics: a review)Microb)Genom)10, 001231 (2024).

26. Hernández Medina, R)et al. Machine learning and deep learning applications in microbiome research)ISME Commun)2, 98 (2022).

27. Przymus, P. et al. Deep learning in microbiome analysis: a comprehensive review of neural network models)Front)Microbiol)15, 1516667 (2024).

28. Marcos-Zambrano, L)J. et al. A toolbox of machine learning software to support microbiome analysis)Front)Microbiol)14, 1250806 (2023).

29. Asnicar, F., Thomas, A)M., Passerini, A., Waldron, L. & Segata, N. Machine learning for microbiologists)Nat)Rev)Microbiol)22, 191–205 (2024).

30. Russakovsky, O. et al. ImageNet large scale visual recognition challenge)Int)J)Comput)Vis)115, 211–252 (2015).

31. Moult, J. A decade of CASP: progress, bottlenecks and prognosis in protein structure prediction)Curr)Opin)Struct)Biol)15, 285–289 (2005).

32. AlQuraishi, M. AlphaFold at CASP13)Bioinformatics 35, 4862–4865 (2019).

33. LaPierre, N., Ju, C)J.-T., Zhou, G. & Wang, W. MetaPheno: A critical evaluation of deep learning and machine learning in metagenome-based disease prediction)Methods 166, 74–82 (2019).

34. Li, M. et al. Performance of gut microbiome as an independent diagnostic tool for 20 diseases: Cross-cohort validation of machine-learning classifiers)Gut Microbes 15, 2205386 (2023).

35. Pasolli, E., Truong, D)T., Malik, F., Waldron, L. & Segata, N. Machine learning meta-analysis of large metagenomic datasets: Tools and biological insights)PLoS Comput)Biol)12, e1004977 (2016).

36. Wang, A. et al. GLUE: A multi-task benchmark and analysis platform for natural language understanding)in Proceedings of the 2018 EMNLP Workshop BlackboxNLP: Analyzing and Interpreting Neural Networks for NLP (eds)Linzen, T., Chrupała, G. & Alishahi, A.) 353–355 (Association for Computational Linguistics, Stroudsburg, PA, USA, 2018).

37. Meyer, F. et al. CAMI Benchmarking Portal: online evaluation and ranking of metagenomic software)Nucleic Acids Res)gkaf369 (2025).

38. Vogtmann, E. et al. Colorectal cancer and the human gut microbiome: Reproducibility with whole-genome shotgun sequencing)PLoS One 11, e0155362 (2016).

39. Yachida, S. et al. Metagenomic and metabolomic analyses reveal distinct stage-specific phenotypes of the gut microbiota in colorectal cancer)Nat)Med)25, 968–976 (2019).

40. Thomas, A)M. et al. Metagenomic analysis of colorectal cancer datasets identifies cross-cohort microbial diagnostic signatures and a link with choline degradation)Nat)Med)25, 667–678 (2019).

41. Feng, Q. et al. Gut microbiome development along the colorectal adenoma-carcinoma sequence)Nat)Commun)6, 6528 (2015).

42. Yu, J. et al. Metagenomic analysis of faecal microbiome as a tool towards targeted non-invasive biomarkers for colorectal cancer)Gut 66, 70–78 (2017).

43. Gupta, A. et al. Association of Flavonifractor plautii, a flavonoid-degrading bacterium, with the gut microbiome of colorectal cancer patients in India)mSystems 4, (2019).

44. Lloyd-Price, J. et al. Multi-omics of the gut microbial ecosystem in inflammatory bowel diseases)Nature 569, 655–662 (2019).

45. Hall, A)B. et al. A novel Ruminococcus gnavus clade enriched in inflammatory bowel disease patients)Genome Med)9, 103 (2017).

46. Li, J. et al. An integrated catalog of reference genes in the human gut microbiome)Nat)Biotechnol)32, 834–841 (2014).

47. Nielsen, H)B. et al. Identification and assembly of genomes and genetic elements in complex metagenomic samples without using reference genomes)Nat)Biotechnol)32, 822–828 (2014).

48. Ijaz, U)Z. et al. The distinct features of microbial ‘dysbiosis’ of Crohn’s disease do not occur to the same extent in their unaffected, genetically-linked kindred)PLoS One 12, e0172605 (2017).

49. Shao, Y. et al. Stunted microbiota and opportunistic pathogen colonization in caesarean-section birth)Nature 574, 117–121 (2019).

50. Bäckhed, F. et al. Dynamics and stabilization of the human gut microbiome during the first year of life)Cell Host Microbe 17, 852 (2015).

51. Vatanen, T. et al. Variation in microbiome LPS immunogenicity contributes to autoimmunity in humans)Cell 165, 1551 (2016).

52. Yassour, M. et al. Strain-level analysis of mother-to-child bacterial transmission during the first few months of life)Cell Host Microbe 24, 146–154.e4 (2018).

53. Wampach, L. et al. Birth mode is associated with earliest strain-conferred gut microbiome functions and immunostimulatory potential)Nat)Commun)9, 5091 (2018).

54. Chu, D)M. et al. Maturation of the infant microbiome community structure and function across multiple body sites and in relation to mode of delivery)Nat)Med)23, 314–326 (2017).

55. Vatanen, T. et al. A distinct clade of Bifidobacterium longum in the gut of Bangladeshi children thrives during weaning)Cell 185, 4280–4297.e12 (2022).

56. Bäckhed, F. et al. Defining a healthy human gut microbiome: current concepts, future directions, and clinical applications)Cell Host Microbe 12, 611–622 (2012).

57. Joos, R. et al. Examining the healthy human microbiome concept)Nat)Rev)Microbiol)23, 192–205 (2025).

58. Human Microbiome Project Consortium)Structure, function and diversity of the healthy human microbiome)Nature 486, 207–214 (2012).

59. Sun, W. et al. A population-scale analysis of 36 gut microbiome studies reveals universal species signatures for common diseases)NPJ Biofilms Microbiomes 10, 96 (2024).

60. Duvallet, C., Gibbons, S)M., Gurry, T., Irizarry, R)A. & Alm, E)J. Meta-analysis of gut microbiome studies identifies disease-specific and shared responses)Nat)Commun)8, 1784 (2017).

61. Zhu, F. et al. Metagenome-wide association of gut microbiome features for schizophrenia)Nat)Commun)11, 1612 (2020).

62. Biau, G. & Scornet, E. A random forest guided tour)Test (Madr.) 25, 197–227 (2016).

63. Chen, T. & Guestrin, C. XGBoost: A Scalable Tree Boosting System)in Proceedings of the 22nd ACM SIGKDD International Conference on Knowledge Discovery and Data Mining (ACM, New York, NY, USA, 2016))doi:10.1145/2939672.2939785.

64. Kleinbaum, D)G. & Klein, M. Logistic Regression: A Self-Learning Text)(Springer, New York, NY, 2010).

65. Suthaharan, S. Support Vector Machine)in Machine Learning Models and Algorithms for Big Data Classification 207–235 (Springer US, Boston, MA, 2016).

66. Austin, G)I. et al. Processing-bias correction with DEBIAS-M improves cross-study generalization of microbiome-based prediction models)Nat)Microbiol)10, 897–911 (2025).

67. Hollmann, N. et al. Accurate predictions on small data with a tabular foundation model)Nature 637, 319–326 (2025).

68. Hollmann, N., Müller, S., Eggensperger, K. & Hutter, F. TabPFN: A Transformer that solves small tabular classification problems in a second)arXiv [cs.LG] (2022) doi:10.48550/arXiv.2207.01848.

69. Murtagh, F. Multilayer perceptrons for classification and regression)Neurocomputing 2, 183–197 (1991).

70. Reiman, D., Metwally, A)A., Sun, J. & Dai, Y. PopPhy-CNN: A Phylogenetic Tree Embedded Architecture for Convolutional Neural Networks to Predict Host Phenotype From Metagenomic Data)IEEE J Biomed Health Inform 24, 2993–3001 (2020).

71. Version 1.6)scikit-learn https://scikit-learn.org/stable/whats_new/v1.6.html.

72. PyTorch documentation — PyTorch 2.6 documentation)https://docs.pytorch.org/docs/2.6/.

73. Pasolli, E. et al. Accessible, curated metagenomic data through ExperimentHub)Nat)Methods 14, 1023–1024 (2017).

