## Supplementary Information for "LAMPP: A benchmark for continuous evaluation of host phenotype prediction from shotgun metagenomic data"

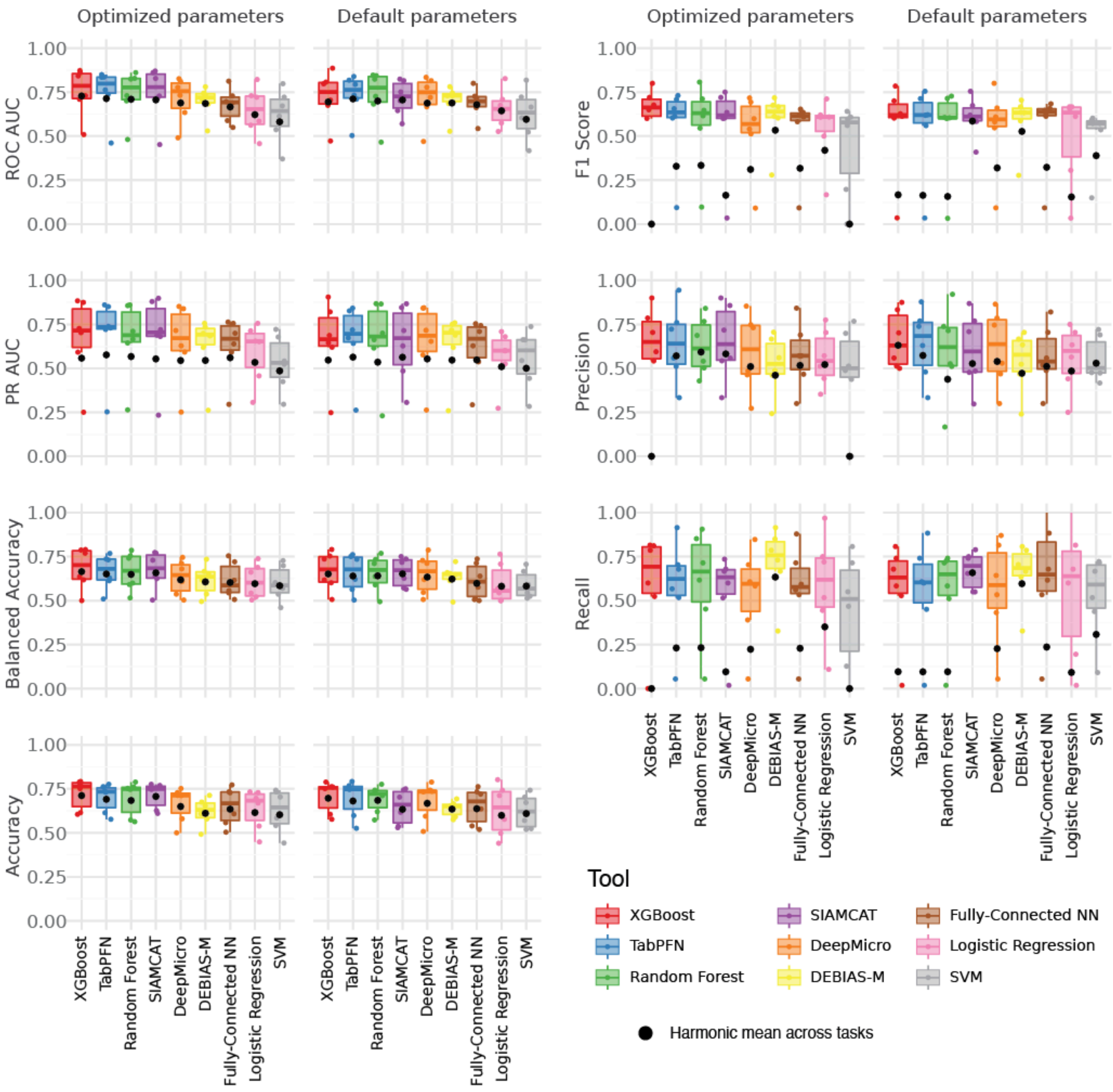


**Fig. S1: Overall performance of baseline tools with optimized and default parameters.** The overall performance of baseline prediction tools with optimized and default parameters over LAMPP's tasks. ROC AUC (**a**), precision-recall (PR) AUC (**b**), balanced accuracy (**c**), accuracy (**d**), F1 score (**e**), precision (**f**), and recall (**g**) of baseline tools (color). Optimized vs default parameters are presented by different facets; Black points mark the harmonic mean across tasks.

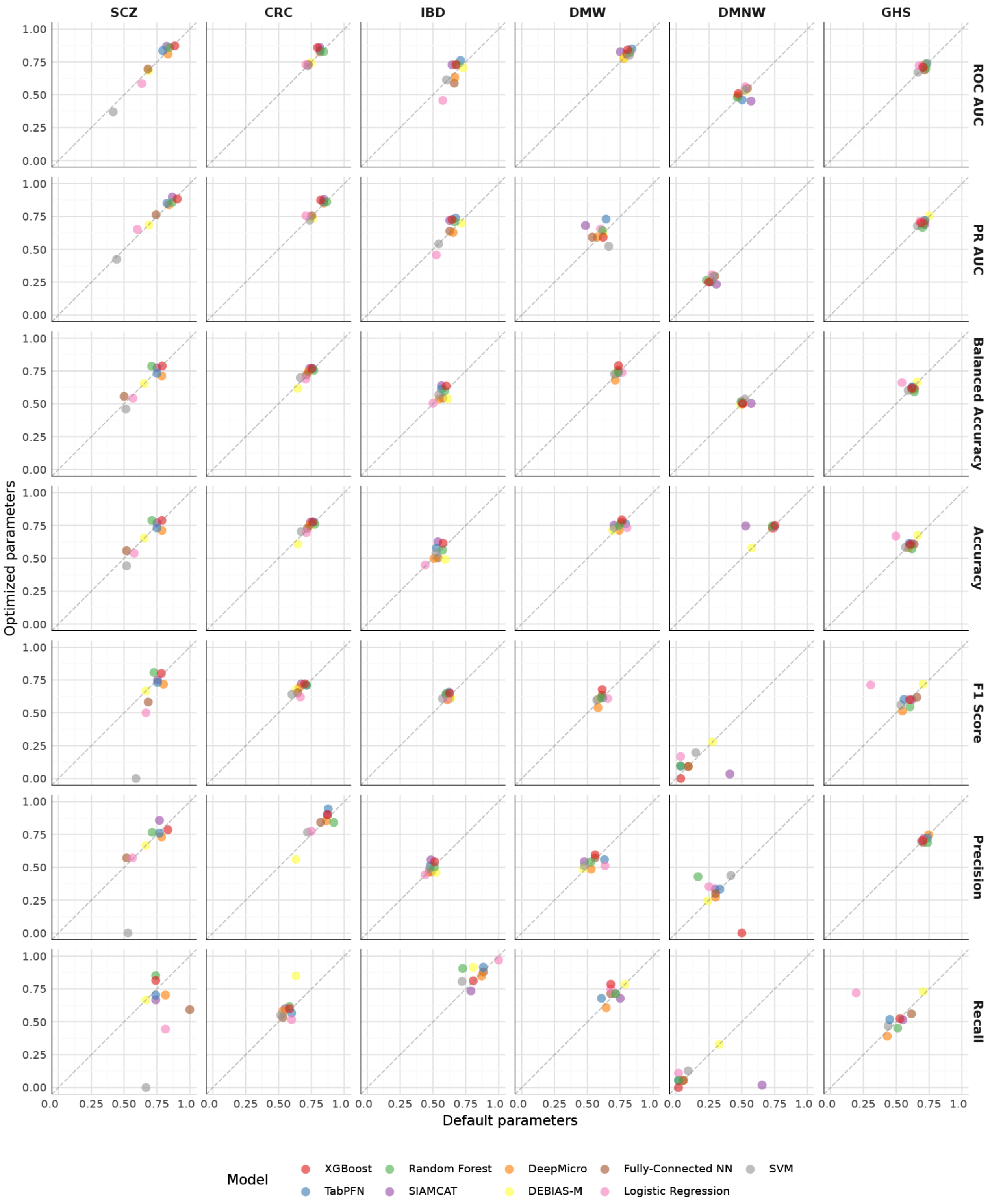


**Fig. S2: Comparison of the performance of baseline tools with optimized vs default parameters.** A comparison of the performance of baseline prediction tools (color) with default (x-axis) and optimized (y-axis) parameters. Plots are faceted by metric (rows) and task (columns). The dashed line marks y=x.

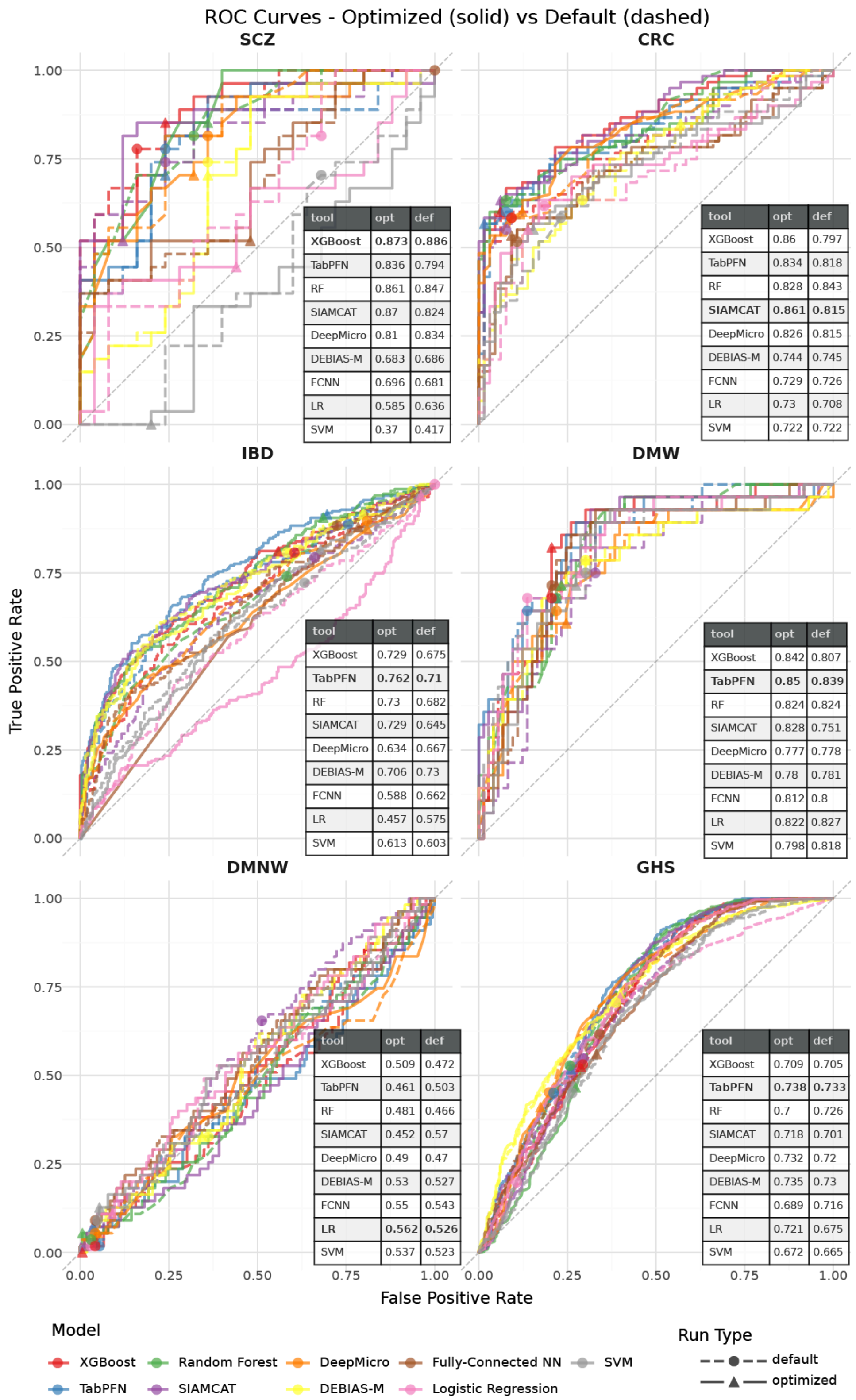


**Fig. S3: ROC curves for baseline tools with optimized and default parameters.** ROC curves for baseline tools with optimized (solid lines, triangle marker) and default (dashed lines, solid circle marker) parameters; The markers indicate the 0.5 decision threshold, and the black diagonal line denotes a random classifier.

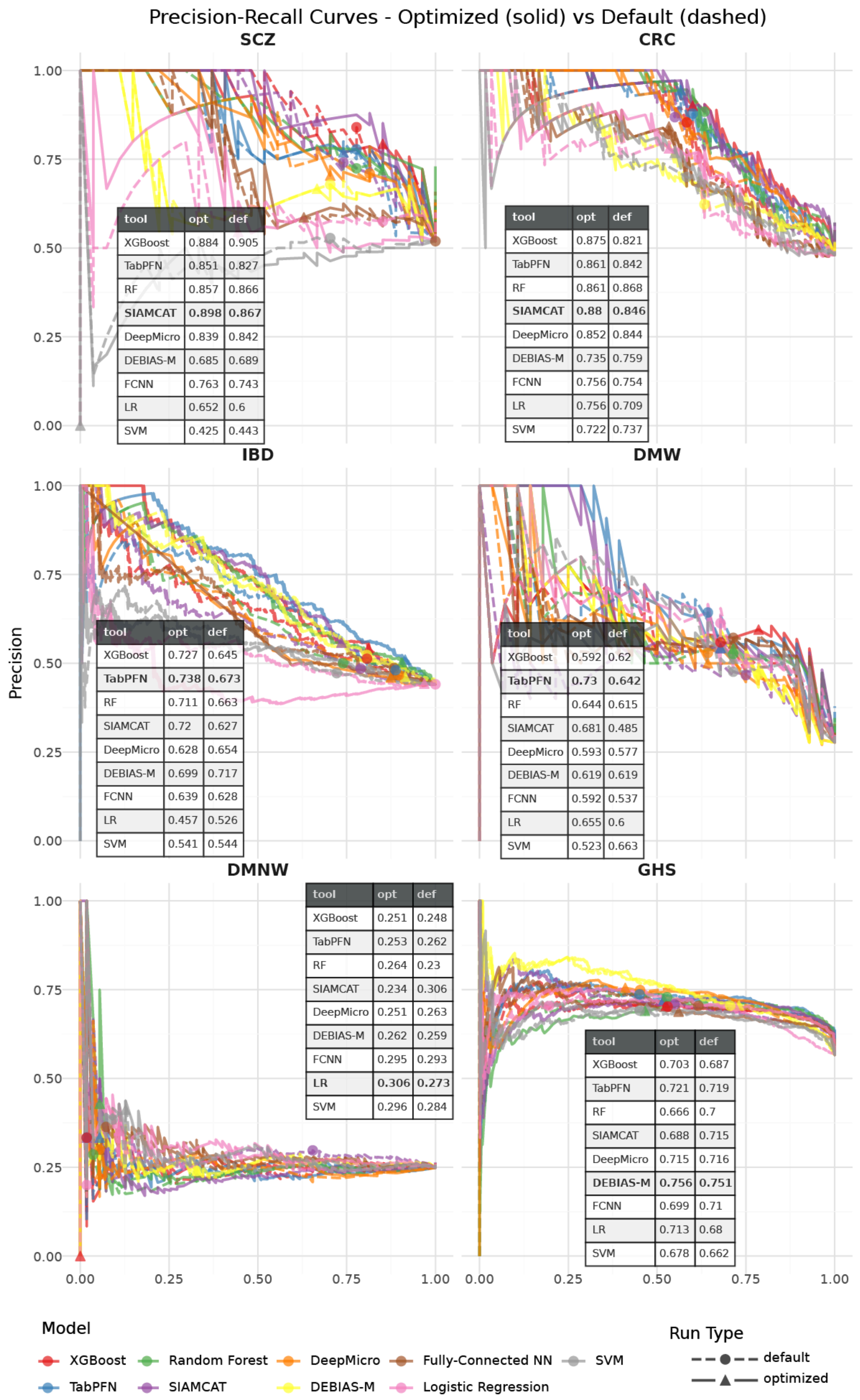


**SCZ**

| tool | opt | def |
|---|---|---|
| XGBoost | 0.884 | 0.905 |
| TabPFN | 0.851 | 0.827 |
| RF | 0.857 | 0.866 |
| **SIAMCAT** | **0.898** | **0.867** |
| DeepMicro | 0.839 | 0.842 |
| DEBIAS-M | 0.685 | 0.689 |
| FCNN | 0.763 | 0.743 |
| LR | 0.652 | 0.6 |
| SVM | 0.425 | 0.443 |

**CRC**

| tool | opt | def |
|---|---|---|
| XGBoost | 0.875 | 0.821 |
| TabPFN | 0.861 | 0.842 |
| RF | 0.861 | 0.868 |
| **SIAMCAT** | **0.88** | **0.846** |
| DeepMicro | 0.852 | 0.844 |
| DEBIAS-M | 0.735 | 0.759 |
| FCNN | 0.756 | 0.754 |
| LR | 0.756 | 0.709 |
| SVM | 0.722 | 0.737 |

**IBD**

| tool | opt | def |
|---|---|---|
| XGBoost | 0.727 | 0.645 |
| **TabPFN** | **0.738** | **0.673** |
| RF | 0.711 | 0.663 |
| SIAMCAT | 0.72 | 0.627 |
| DeepMicro | 0.628 | 0.654 |
| DEBIAS-M | 0.699 | 0.717 |
| FCNN | 0.639 | 0.628 |
| LR | 0.457 | 0.526 |
| SVM | 0.541 | 0.544 |

**DMW**

| tool | opt | def |
|---|---|---|
| XGBoost | 0.592 | 0.62 |
| **TabPFN** | **0.73** | **0.642** |
| RF | 0.644 | 0.615 |
| SIAMCAT | 0.681 | 0.485 |
| DeepMicro | 0.593 | 0.577 |
| DEBIAS-M | 0.619 | 0.619 |
| FCNN | 0.592 | 0.537 |
| LR | 0.655 | 0.6 |
| SVM | 0.523 | 0.663 |

**DMNW**

| tool | opt | def |
|---|---|---|
| XGBoost | 0.251 | 0.248 |
| TabPFN | 0.253 | 0.262 |
| RF | 0.264 | 0.23 |
| SIAMCAT | 0.234 | 0.306 |
| DeepMicro | 0.251 | 0.263 |
| DEBIAS-M | 0.262 | 0.259 |
| FCNN | 0.295 | 0.293 |
| **LR** | **0.306** | **0.273** |
| SVM | 0.296 | 0.284 |

**GHS**

| tool | opt | def |
|---|---|---|
| XGBoost | 0.703 | 0.687 |
| TabPFN | 0.721 | 0.719 |
| RF | 0.666 | 0.7 |
| SIAMCAT | 0.688 | 0.715 |
| DeepMicro | 0.715 | 0.716 |
| **DEBIAS-M** | **0.756** | **0.751** |
| FCNN | 0.699 | 0.71 |
| LR | 0.713 | 0.68 |
| SVM | 0.678 | 0.662 |

**Fig. S4: Precision-Recall curves for baseline tools with optimized and default parameters.** ROC curves for baseline tools with optimized (solid lines, triangle marker) and default (dashed lines, solid circle marker) parameters; The markers indicate the 0.5 decision threshold.

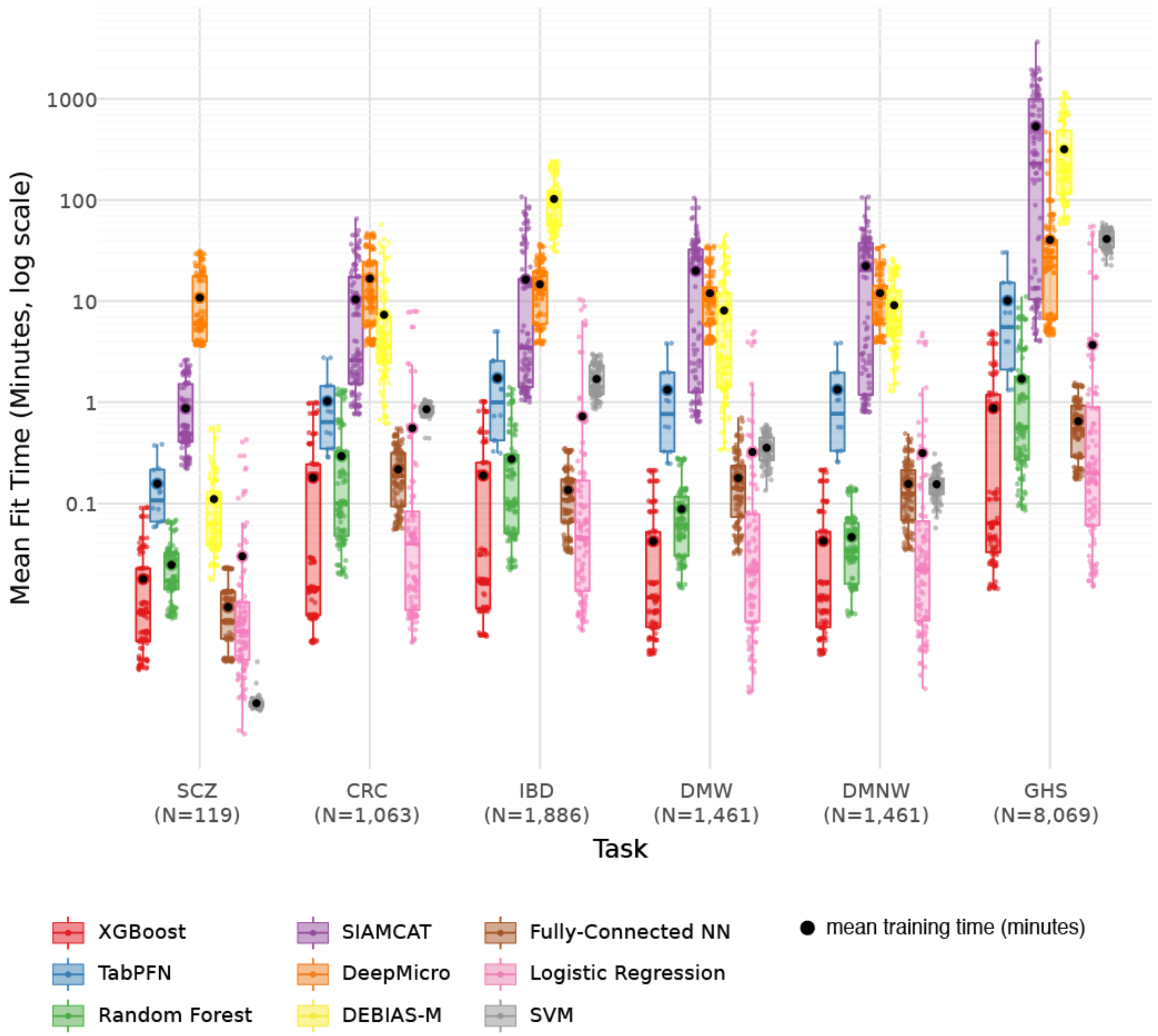


**Fig. S5: Training time of baseline tools across parameter configurations.** Total training time for 5-fold cross-validation configurations by tool (color) and task. The average training time is marked in black on each boxplot.

### Methods S1: Parameter configurations tested

Here are all the parameter configurations tested for each tool. Default parameters are shown in **bold**.

**XGBoost**

1. n_estimators (number of boosting stages): 50, **100**, 200
2. learning_rate (boosting learning rate): 0.05, **0.1**, 0.5
3. loss (loss function): **log_loss**, exponential
4. max_features (features per split): sqrt, log2, **None**
5. subsample (fraction of samples per tree): 0.8, **1.0**

**Total of 108 configurations**

**TabPFN**

1. n_estimators: 1, 2, 4, **8**, 16, 32
2. balance_probabilities: True, **False**
3. Ignore_pretraining_limits: True

**Total of 12 configurations**

**Random Forest**

1. n_estimators (number of trees): 50, **100**, 200
2. max_depth (maximum tree depth): **None**, 20
3. criterion (split quality measure): **gini**, entropy, log_loss
4. max_features (features per split): **sqrt**, log2, 0.2
5. class_weight (class weight adjustment): balanced, **None**

**Total of 108 configurations**

**SIAMCAT**

1. cutoff (cutoff for data filtering): **0.001**, 0.005, 0.01
2. norm.method (data normalization method): rank.unit, rank.std, **log.std**, log.unit, log.clr, std
3. method (machine learning classifier): **lasso**, enet, ridge, lasso_ll, ridge_ll, randomForest

**Total of 108 configurations**

* For the GHS task, SIAMCAT runs with cutoff=0.01 failed with the error "No features retained after filtering! Try changing your cutoff. Exiting…" so only 72 configurations were tested.

**DeepMicro**

1. Autoencoder architectures:
   a. AE dims (Autoencoder layer dimensions): [64], [128], [256], [64,32], [128,64], [256,128]
   b. VAE dims (Variational Autoencoder layer dimensions): [8], [16], [32], [16,8], [32,16], [64,32]
   c. **CAE** dims (Convolutional Autoencoder layer dimensions): **[32]**, [16], [32,16,8], [16,8,4], [16,8], [8,4]
2. Classifier configurations:
   a. **RF** (Random Forest) criterion (split quality measure): **gini**, entropy
   b. SVM (Support Vector Machine) kernel (kernel type): linear, rbf
   c. MLP (Multi-Layer Perceptron) numHiddenLayers (number of hidden layers): 1, 2

**Total of 108 configurations** (18 autoencoder configurations × 6 classifier configurations)

**DEBIAS-M**

1. learning_rate: 0.0005, 0.001, **0.005**, 0.01
2. Min_epochs (the minimum number of training epochs): 10, **25**, 40
3. l2_strength (the l2 regularization of the linear predictive layer's parameters. Larger values specify stronger regularization): **0**, 0.01, 0.1
4. W_l2 (The l2 regularization of the multiplicative bias correction parameters. Larger values specify stronger regularization): **0**, 0.01, 0.1

**Total of 108 configurations**

**Fully Connected Neural Network**

1. hidden_dim (hidden layer dimension): 16, 32, **64**, 128
2. lr (learning rate): 0.01, **0.001**, 0.0001
3. epochs (number of training epochs): **20**, 50, 100
4. batch_size (training batch size): 64, **128**, 256

**Total of 108 configurations**

**Logistic Regression**

1. penalty (regularization type): **l2**, l1, elasticnet, None
2. solver (optimization algorithm): **lbfgs**, saga, newton-cg, sag, liblinear
3. C (inverse regularization strength): 0.01, 0.1, **1**, 10, 100
4. l1_ratio (elastic net mixing parameter): 0.1, 0.5, 0.9 (for elasticnet only)
5. class_weight (class weight adjustment): **None**, balanced

**Total of 108 configurations***

* Different penalties accept different parameters, so out of the 600 potential configurations, only 108 were valid: 50 for L2, 20 for l1, 30 for elasticnet and 8 with None regulation.

**SVM**

1. kernel (kernel type): linear, **rbf**, poly, sigmoid
2. C (regularization parameter): 0.01, 0.1, **1**, 10
3. gamma (kernel coefficient): **scale**, auto, 0.001
4. class_weight (class weight adjustment): balanced, **None**

**Total of 96 configurations**

### Methods S2: Top-performing configurations by tool and task

**XGBoost**

| task | max_features | n_estimators | learning_rate | loss | subsample | AUC averaged across folds |
|---|---|---|---|---|---|---|
| **SCZ** | sqrt | 200 | 0.05 | exponential | 0.8 | 0.884 |
| **CRC** | sqrt | 200 | 0.05 | log_loss | 1 | 0.798 |
| **DMW** | sqrt | 200 | 0.05 | exponential | 0.8 | 0.888 |
| **DMNW** | sqrt | 200 | 0.05 | exponential | 0.8 | 0.888 |
| **IBD** | log2 | 200 | 0.05 | exponential | 0.8 | 0.833 |
| **GHS** | None | 200 | 0.1 | log_loss | 0.8 | 0.893 |

**TabPFN**

| task | n_estimators | balance_probabilities | AUC averaged across folds |
|---|---|---|---|
| **SCZ** | 32 | TRUE | 0.846 |
| **CRC** | 32 | TRUE | 0.802 |
| **DMW** | 32 | TRUE | 0.881 |
| **DMNW** | 32 | TRUE | 0.881 |
| **IBD** | 32 | TRUE | 0.746 |
| **GHS** | 32 | TRUE | 0.883 |

**Random Forest**

| task | class_weight | criterion | max_depth | max_features | n_estimators | AUC averaged across folds |
|---|---|---|---|---|---|---|
| **SCZ** | None | gini | None | sqrt | 50 | 0.859 |
| **CRC** | balanced | entropy | 20 | 0.2 | 50 | 0.806 |
| **DMW** | balanced | entropy | 20 | sqrt | 200 | 0.891 |
| **DMNW** | balanced | entropy | 20 | sqrt | 200 | 0.891 |
| **IBD** | balanced | gini | 20 | log2 | 200 | 0.805 |

| | | | | | | |
|---|---|---|---|---|---|---|
| **GHS** | balanced | entropy | None | 0.2 | 200 | 0.896 |

**SIAMCAT**

| task | filter_cutoff | norm | ml | AUC averaged across folds |
|---|---|---|---|---|
| **SCZ** | 0.01 | log.std | ridge_ll | 0.877 |
| **CRC** | 0.005 | std | randomForest | 0.808 |
| **DMW** | 0.001 | log.std | randomForest | 0.886 |
| **DMNW** | 0.005 | log.unit | randomForest | 0.888 |
| **IBD** | 0.005 | rank.std | ridge_ll | 0.788 |
| **GHS** | 0.001 | log.unit | randomForest | 0.88 |

**DeepMicro**

| task | ae_dims | autoencoder | classifier | clf_criterion | AUC averaged across folds |
|---|---|---|---|---|---|
| **SCZ** | [32] | cae | rf | gini | 0.825 |
| **CRC** | [16] | cae | rf | entropy | 0.768 |
| **DMW** | [16, 8] | cae | rf | entropy | 0.876 |
| **DMNW** | [16] | cae | rf | entropy | 0.878 |
| **IBD** | [32] | cae | rf | entropy | 0.775 |
| **GHS** | [32] | cae | rf | entropy | 0.882 |

**DEBIAS-M**

| task | learning_rate | min_epochs | l2_strength | w_l2 | batch_str | AUC averaged across folds |
|---|---|---|---|---|---|---|
| **SCZ** | 0.01 | 40 | 0 | 0 | 0 | 0.72 |
| **CRC** | 0.01 | 40 | 0 | 0.1 | infer | 0.695 |
| **DMW** | 0.005 | 40 | 0 | 0 | infer | 0.862 |
| **DMNW** | 0.005 | 25 | 0 | 0 | infer | 0.861 |
| **IBD** | 0.001 | 10 | 0 | 0 | infer | 0.721 |
| **GHS** | 0.001 | 40 | 0 | 0 | infer | 0.953 |

**Fully Connected Neural Network**

| task | hidden_dim | lr | epochs | batch_size | AUC averaged across folds |
|---|---|---|---|---|---|
| **SCZ** | 128 | 0.01 | 20 | 64 | 0.697 |
| **CRC** | 32 | 0.001 | 20 | 64 | 0.703 |
| **DMW** | 16 | 0.001 | 20 | 64 | 0.853 |
| **DMNW** | 16 | 0.001 | 20 | 64 | 0.853 |
| **IBD** | 16 | 0.01 | 100 | 256 | 0.692 |
| **GHS** | 128 | 0.001 | 20 | 64 | 0.843 |

**Logistic Regression**

| task | C | class_weight | penalty | solver | AUC averaged across folds |
|---|---|---|---|---|---|
| **SCZ** | 100 | None | l1 | liblinear | 0.655 |
| **CRC** | 100 | balanced | l2 | liblinear | 0.663 |
| **DMW** | 10 | balanced | l2 | newton-cg | 0.848 |
| **DMNW** | 10 | balanced | l2 | newton-cg | 0.848 |
| **IBD** | 1 | None | l1 | saga | 0.646 |
| **GHS** | 100 | balanced | l1 | liblinear | 0.833 |

**SVM**

| task | C | class_weight | gamma | kernel | AUC averaged across folds |
|---|---|---|---|---|---|
| **SCZ** | 0.01 | None | scale | poly | 0.591 |
| **CRC** | 10 | balanced | scale | linear | 0.659 |
| **DMW** | 10 | None | scale | linear | 0.848 |
| **DMNW** | 10 | None | scale | linear | 0.848 |
| **IBD** | 10 | None | scale | rbf | 0.641 |
| **GHS** | 10 | None | scale | rbf | 0.815 |
